# GPU-accelerated alignment of bisulfite-treated short-read sequences

**DOI:** 10.1101/175729

**Authors:** Richard Wilton, Xin Li, Andrew P. Feinberg, Alexander S. Szalay

## Abstract

The alignment of bisulfite-treated DNA sequences (BS-seq reads) to a large genome involves a significant computational burden beyond that required to align non-bisulfite-treated reads. In the analysis of BS-seq data, this can present an important performance bottleneck that can potentially be addressed by appropriate software-engineering and algorithmic improvements. One strategy is to integrate this additional programming logic into the read-alignment implementation in a way that the software becomes amenable to optimizations that lead to both higher speed and greater sensitivity than can be achieved without this integration.

We have evaluated this approach using Arioc, a short-read aligner that uses GPU (general-purpose graphics processing unit) hardware to accelerate computationally-expensive programming logic. We integrated the BS-seq computational logic into both GPU and CPU code throughout the Arioc implementation. We then carried out a read-by-read comparison of Arioc's reported alignments with the alignments reported by the most widely used BS-seq read aligners. With simulated reads, Arioc's accuracy is equal to or better than the other read aligners we evaluated. With human sequencing reads, Arioc's throughput is at least 10 times faster than existing BS-seq aligners across a wide range of sensitivity settings.

The Arioc software is available at https://github.com/RWilton/Arioc. It is released under a BSD open-source license.

## INTRODUCTION

As the use of next-generation DNA sequencing technology becomes increasingly widespread, the cost of sequencing a single human genome at 30-fold coverage continues to decrease toward $1,000 (van Nimwegen et al., 2016), and the number of large datasets generated by next-generation sequencing is growing. The first step in analyzing the data generated by a DNA sequencing run is read alignment, the process of determining the point of origin of each sequencing read with respect to a reference genome. Read alignment is algorithmically complex and time consuming, to the point where the time spent in executing read-alignment software approaches that of the DNA sequencing run itself.

To address this need, a number of attempts have been made to develop read-alignment software that exploits the parallel processing capability of general-purpose graphics processing units, or GPUs (Schatz et al., 2007). GPUs are video display devices whose hardware and system-software architecture can also be used for general purpose computing. They are well suited to software implementations where independent computations on many thousands of data items can be carried out in parallel. This was the primary motivation for the development of Arioc (Wilton et al., 2015), a high-throughput GPU-based read aligner.

There are additional computational challenges in aligning DNA sequencing reads when the DNA has been treated with bisulfite so as to differentiate methylcytosine from cytosine residues in the DNA sequences. After bisulfite-treated DNA is sequenced, the resulting BS-seq short reads must be aligned to a reference genome in a manner that identifies each methylcytosine occurrence in the context of its neighboring bases. Arioc extracts this information from BS-seq reads using software techniques that provide for efficient GPU acceleration.

### Encoding of methylcytosine bases in BS-seq reads

Bisulfite treatment of DNA converts cytosine residues to uracil while leaving methylcytosine residues intact; uracil is subsequently replaced by thymine during PCR amplification of the DNA. Thus, in effect, thymine in the sequencer reads for bisulfite-treated DNA can represent either thymine or methylcytosine in the original DNA:

| <u>In untreated DNA</u> | <u>In sequencer reads</u> |
| --- | --- |
| A | A |
| C<sub>m</sub> (methylcytosine) | C |
| G | G |
| C or T | T |

The accepted technique for disambiguating the Ts in a BS-seq read sequence is to compare the read sequence to the corresponding reference sequence at the location where the read is properly aligned to the reference. Each T in the read may then be inferred to map to the reference as follows:

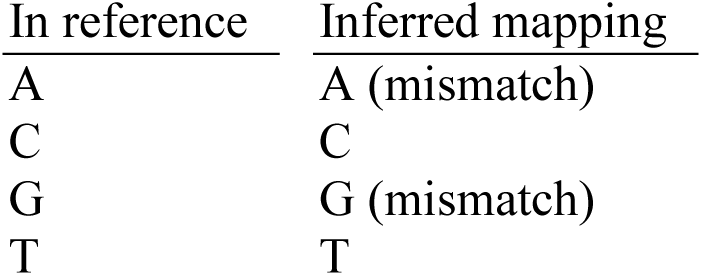

| <u>In reference</u> | <u>Inferred mapping</u> |
| --- | --- |
| A | A (mismatch) |
| C | C |
| G | G (mismatch) |
| T | T |

This CT ambiguity introduces additional complexity into the sequence-alignment procedure in two significant ways:

- Where a read aligner uses lookup tables, indexes, or other data structures to identify reference locations at which potential alignments may be found for a read, those data structures must be constructed so as to accommodate ambiguous Ts.
- The read aligner must consider the inferred mapping of ambiguous Ts to the reference sequence when it computes a numeric alignment score for a mapped read.

This ambiguity in the encoding of BS-seq reads increases the complexity of the software implementation of the read aligner. It represents an additional computational burden that decreases the overall speed of the software in comparison with the alignment process for non-bisulfite-treated reads. In Arioc, the programming logic involved in handling BS-seq reads is integrated into the general-purpose read-alignment implementation so that the entire software pipeline remains highly parallel and amenable to GPU acceleration.

## METHODS

The Arioc aligner is written in C++ and compiled for both Windows (with Microsoft Visual C++) and Linux (with the GNU C++ compiler). The implementation runs in a single computer on a user-configurable number of concurrent CPU threads and on one or more NVidia GPUs. The implementation pipeline uses 38 different CUDA kernels written in C++ (nongapped and gapped alignment computation, application-specific list processing) and about 150 calls to various CUDA Thrust APIs (sort, set reduction, set difference, string compaction).

We used two development and test computers for BS-seq experiments with Arioc, one with Microsoft Windows Server 2008 R2 and one with RedHat Scientific Linux release 7.3. Each computer was built with dual 6-core Intel Xeon X5670 CPUs running at 2.93GHz, so 24 logical threads were available to applications. There was 144GB of system RAM, of which about 96GB was available to applications. Each computer was also configured with three NVidia Tesla series GPUs (Kepler K20c), each of which supports 5GB of on-device "global" memory and 26624 parallel threads. The internal expansion bus in each machine was PCIe v2.

Throughput (query sequences aligned per second) was measured when the test computers were otherwise idle so that all CPU, memory, and I/O resources were available. For experiments with simulated data, we used Sherman (Krueger, 2014) to generate 100 nt paired-end reads. For experiments with Illumina data, we used 100 nt paired-end Illumina HiSeq 2500 BS-seq data from an "Omics catalogue of lung adenocarcinoma cell lines" (Suzuki et al., 2014).

### Software implementation

As described elsewhere (Wilton et al., 2015), Arioc is implemented as a pipeline in which batches of reads are processed by a sequence of discrete software modules, each of which operates on a separate CPU thread that is allocated for the lifetime of the module and then discarded. When multiple GPUs are used, each GPU is associated with its own CPU thread. Modules execute concurrently on CPU threads and on the GPU.

Arioc aligns reads by first extracting short subsequences (seeds) from each read. It then uses lookup (hash) tables to identify reference-sequence locations at which each seed subsequence appears in the reference sequence (genome). Specific adaptations within the Arioc implementation for aligning BS-seq reads are:

- Seed lookup tables in which all Cs are represented by Ts.
- Disambiguation of Ts in mapped read sequences by comparison with the reference sequence.
- Read-alignment scoring based on disambiguated read-sequence mapping.
- Creation of a methylation context map for each mapped read.

For each bisulfite-treated read, Arioc converts Cs to Ts in the read sequence. It then uses its CT-converted lookup tables to find high-priority reference-sequence locations at which to carry out alignments. It computes and scores alignments by performing a base-by-base comparison of the original bisulfite-treated read with the original (not CT-converted) reference sequence.

### Read alignment

Arioc performs read alignment in two passes. It first attempts nongapped spaced-seed alignment (Chen et al., 2009) in order to quickly identify read sequences that differ from the reference by no more than a few mismatches, without insertions or deletions. If the candidate location was identified using the reverse complement of the read sequence, the alignment is done using that reverse complement. Each alignment is scored by allowing a T in the read sequence to match either C or T in the reference.

For reads that do not have a sufficient number of nongapped mappings, Arioc computes base-by-base alignments at each candidate location using the Smith-Waterman algorithm (Smith and Waterman, 1981). The computation is again carried out between the original reference and read sequences. Again, Arioc computes alignment scores with either C or T in the reference sequence matching T in the read sequence. If the candidate location was identified by seeding the reverse complement of the read sequence, the alignment is done using that reverse complement.

This technique ensures that the base mapping (as reported in the SAM CIGAR field) and alignment score are accurately computed, regardless of the presence of methylcytosine in the read sequence. Furthermore, when aligning paired-end reads, Arioc applies the same heuristics for identifying high-priority candidate locations as it does when aligning non-bisulfite-treated reads. For example, Arioc prioritizes pairs of candidate locations that are consistent with user-specified orientation and fragment-length limits.

### Managing the CT ambiguity

Like most other BS-seq aligners, Arioc deals with CT ambiguity using lookup tables (LUTs) or indexes in which all Cs are represented as Ts. Arioc probes the LUTs or indexes using seeds (subsequences of each bisulfite-treated read) in which all Cs are likewise replaced by Ts (Figure 1). Arioc also probes the LUTs using seeds derived from the CT-converted reverse complement of the read sequence.

**Figure 1.**
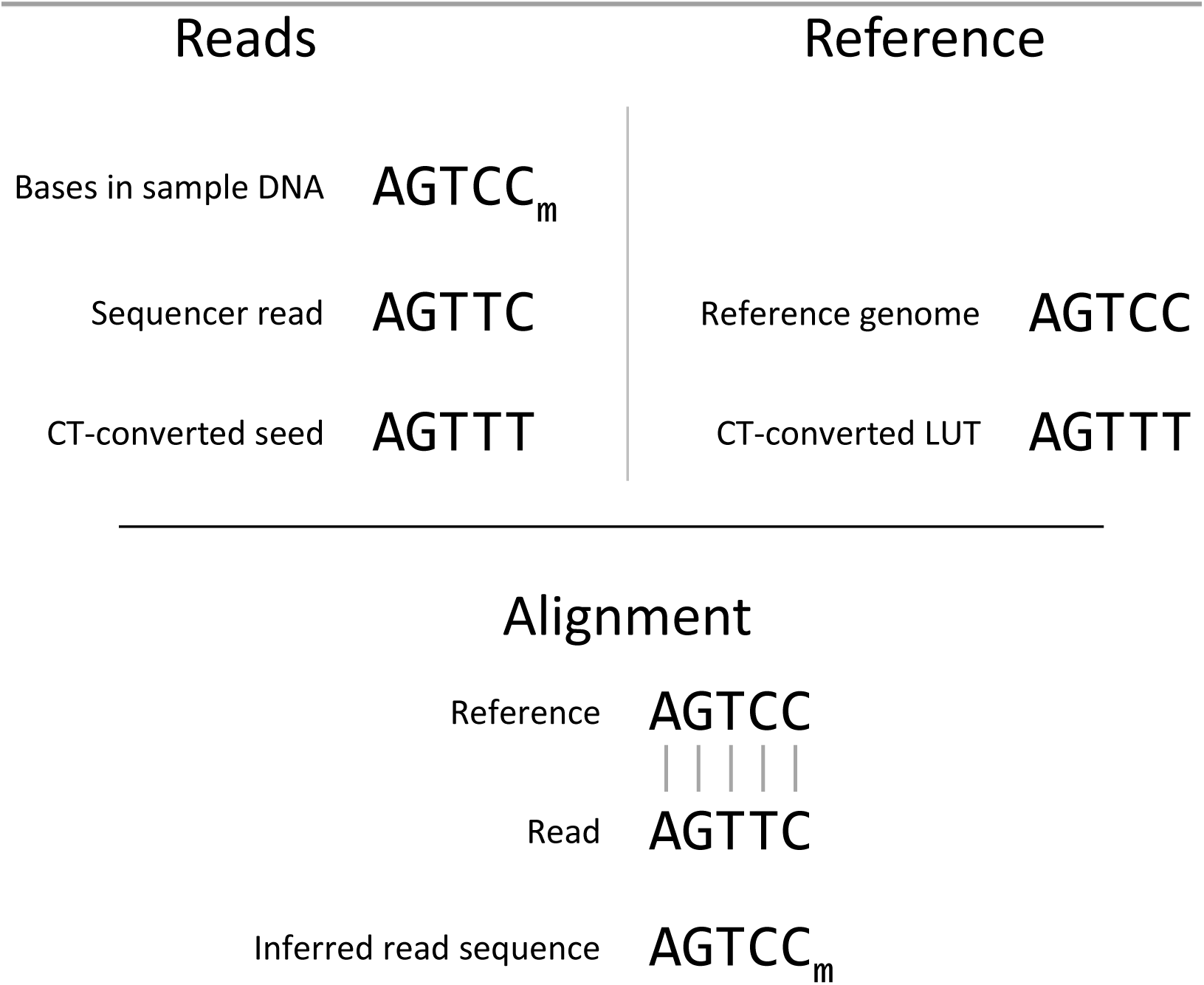
Bisulfite treatment of a DNA sample that contains both cytosine and methylcytosine (C_m_) results in sequencer reads that contain thymine in positions where the original DNA contains cytosine and cytosine in positions where the original DNA contains methylcytosine. Arioc converts all Cs to Ts in each sequencer read and probes a CT-converted lookup table for reference-genome locations at which to compute alignments between the original reference-genome sequence and the read sequence.

The result is a list of candidate locations at which to perform base-by-base alignment between the reference and read sequences. Arioc sorts the candidate-locations list and, for each read and candidate location, counts the number of seeds that reference each location. Candidates are then prioritized for alignment based on the number of different seeds that fall within a sufficiently narrow range of adjacent locations. All of these operations are well suited to GPU acceleration.

### Identifying methylation sites

Once Arioc has identified the set of high-scoring mappings to report, it performs a base-by-base comparison of each mapped read sequence with the corresponding region in the reference sequence (genome). This procedure follows that implemented in Bismark (Krueger and Andrews, 2011): Arioc establishes a methylation context (CpG, CHG, CHH, CHN, CN) for each methylcytosine by examining the two subsequent bases in the read sequence. It reports the position and methylation context of each identified methylcytosine in a character-string map associated with the read sequence (emitted as an optional XM field in SAM-formatted alignment results).

### Analysis of alignment results

We used the human reference genome release 38 (Genome Reference Consortium, 2016) for throughput and sensitivity experiments. We evaluated published speed and sensitivity results for a number of BS-seq aligners (Supplementary Table T1) and identified three programs whose reported speed and sensitivity with BS-seq reads made them acceptable candidates for direct comparison with the Arioc implementation:

- Bismark (CPU)
- Segemehl (Otto et al., 2012) (CPU)
- GPU-BSM (Manconi et al., 2014) (GPU)

We parsed the SAM-formatted output (SAM/BAM Format Specification Working Group, 2016) from each aligner and aggregated the alignments reported by each aligner for each read. We examined the POS (position), TLEN (paired-end fragment length), and AS (alignment score) fields to compare the mappings reported for each read by each aligner. We computed local alignments using the following scoring parameters: match=+2; mismatch=−6; gap open=−5; gap space=−3, with a threshold alignment score of one half of the maximum possible alignment score.

We used simulated (Sherman) reads to evaluate sensitivity for both paired-end and unpaired reads. For each aligner, we used high "effort" parameters so as to favor sensitivity over throughput. We assumed that a read was correctly mapped when, after accounting for soft clipping, one or both of its ends mapped within 40 nt of the mapping generated by Sherman. (Supplementary Table T4 explains our choice of a 40 nt threshold.) To illustrate sensitivity and specificity, we plotted the cumulative number of correctly-mapped and incorrectly-mapped reads reported by each aligner, stratified by the MAPQ score (Li, Ruan, and Durbin, 2008) for each read.

We used the Illumina BS-seq lung adenocarcinoma data to measure throughput using both paired-end and unpaired reads. For this analysis, we recorded throughput across a range of parameters chosen so as to trade speed for sensitivity. We defined "sensitivity" as the percentage of reads reported as mapped by each aligner with alignment score (and, for paired-end reads, TLEN) within configured limits.

Prior to computing alignments, the GPU-aware aligners spend a brief period of execution time initializing static data structures in GPU device memory. We excluded this startup time from throughput calculations for these aligners.

## RESULTS

Each of the read aligners we tested is able to map tens of millions of reads to the human genome in an acceptably short period of time. Each was capable of mapping simulated reads with high accuracy. With sequencer reads, Arioc demonstrated up to 10 times higher throughput across a wide range of sensitivity settings.

### Evaluation with simulated reads

With simulated Illumina read data, Arioc was able to map paired-end reads to their correct origin in the reference genome with sensitivity and specificity as good as or better than each of the other aligners to which we compared it (Figure 2 and Supplementary Figures S1-S8). Arioc maintains a very high ratio of correct to incorrect mappings until mappings with relatively low MAPQ scores are considered.

**Figure 2.**
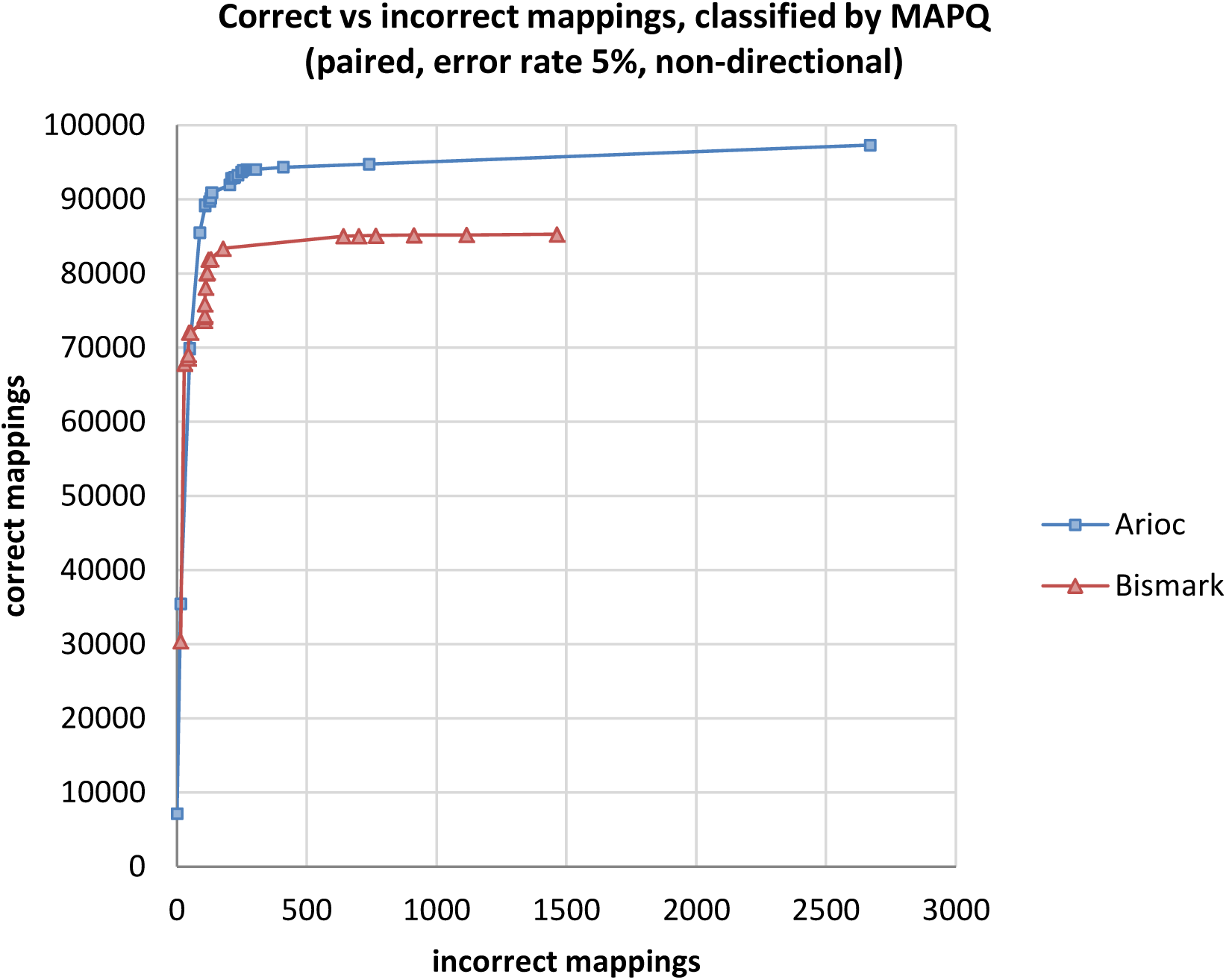
Total correctly mapped versus incorrectly mapped reads, plotted for decreasing MAPQ, for 100,000 simulated 100nt paired-end Illumina reads (200,000 mates).

### Evaluation with sequencer-generated reads

We used the lung adenocarcinoma data to evaluate speed (Figure 3). Across a wide range of sensitivity settings, Arioc's speed when executed across three GPUs is over 10 times that of the CPU-based aligners to which we compared it, and over five times that of the GPU-based aligner to which we compared it. Furthermore, Arioc's throughput scales almost linearly when two or more GPUs are available (Supplementary Figure S10).

**Figure 3.**
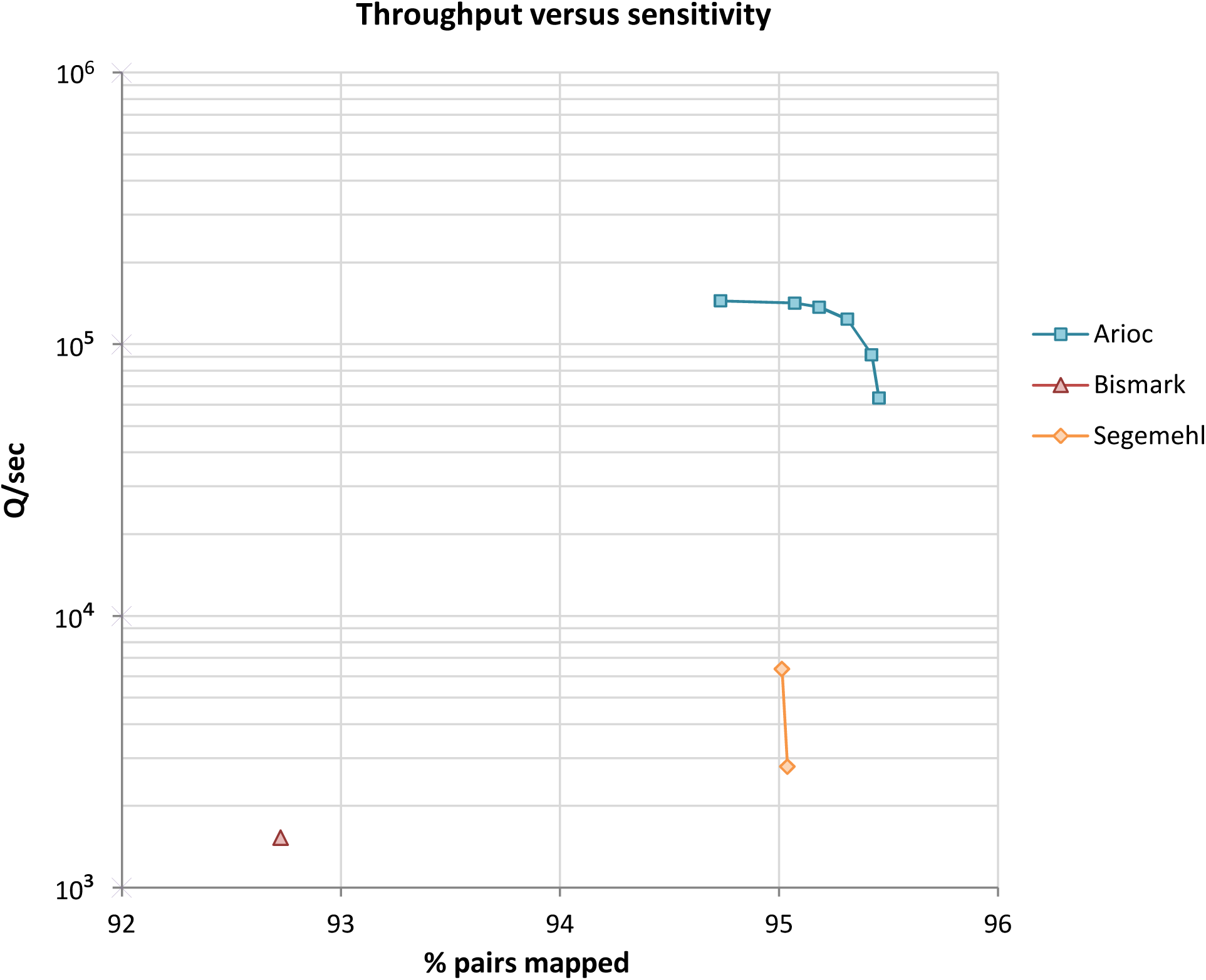
Speed (reads aligned per second) versus sensitivity (percentage of paired-end reads mapped) for three BS-seq aligners. Data for 4 million 100 nt paired-end BS-seq reads (Suzuki et al., 2014). Workstation hardware: 12 CPU cores (24 threads of execution), one NVidia K20c GPU.

## DISCUSSION

The Arioc implementation demonstrates that high throughput can be achieved by GPU acceleration without losing sensitivity. Furthermore, by sacrificing throughput, Arioc can be "pushed" to achieve greater sensitivity.

### Lookup tables and seeds for BS-seq short-read alignment

Arioc uses lookup tables to identify the set of reference-sequence locations that correspond to each seed it derives from a read sequence. For each possible seed, the lookup table contains a list of reference locations that correspond to a bitwise numerical hash of the seed sequence, that is, the lookup table is a simple hash table whose bins each contain the reference-sequence locations for one seed hash value.

Choosing an appropriate seed length is important to achieving greater processing speed. As seed length is increased the average hash bin size decreases, so the read aligner performs alignments at fewer reference-sequence locations and overall throughput improves. Furthermore, longer seeds are intrinsically more specific for highly-similar reference-sequence locations. As seeds become longer, however, they are increasingly likely to span mismatches, insertions, and deletions within read sequences and thus to fail to identify the best reference-sequence locations at which to perform alignments.

With non-CT-converted short reads, a seed length of 20 nt provides a reasonable balance between hash efficiency and sensitivity; for this reason, 20 nt is the default seed length in both Bowtie 2 and Arioc. With CT-converted DNA, however, a seed length of about 25 nt may be used in order to obtain a similar balance.

**Figure 4.**
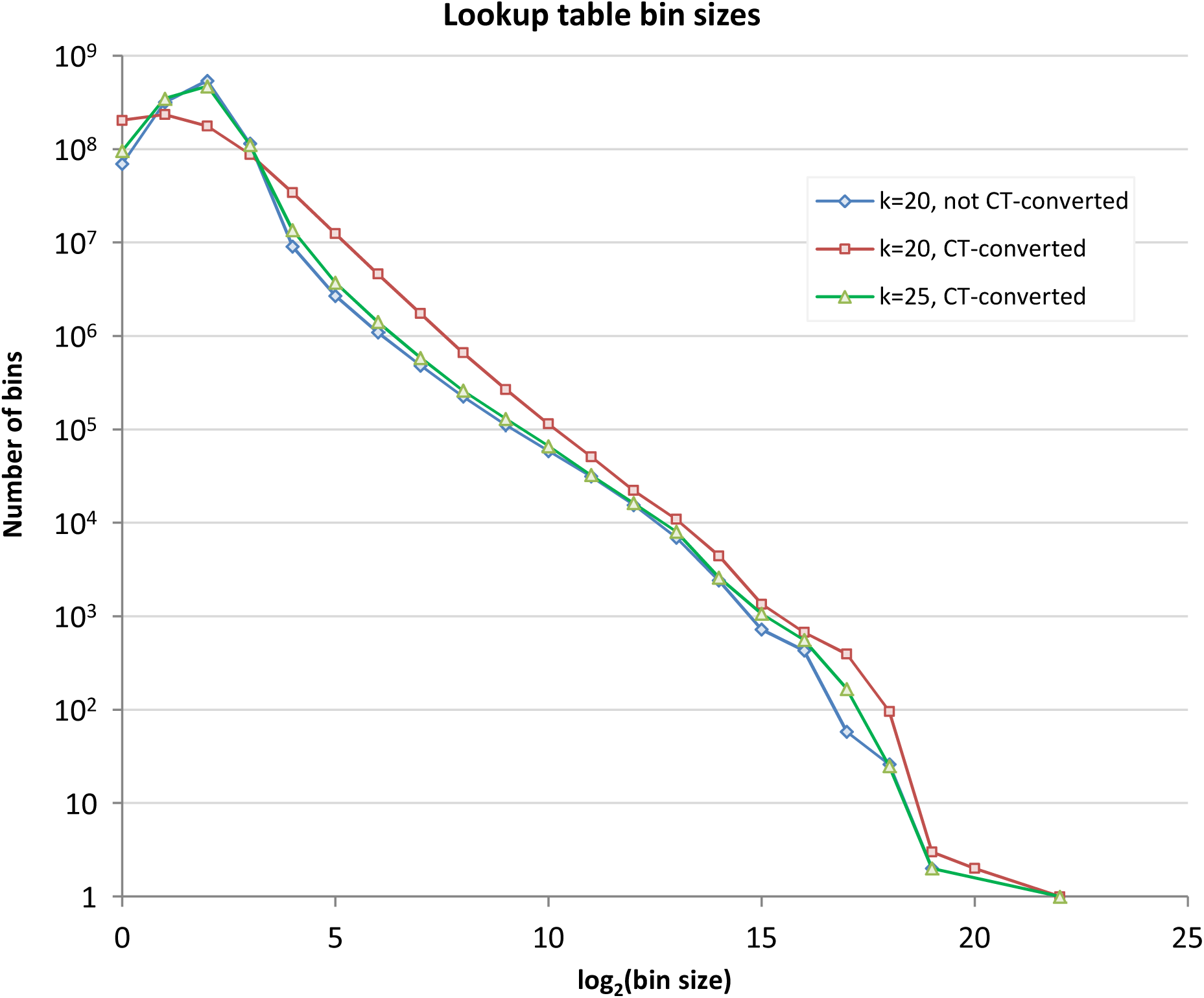
Lookup table bin sizes for CT-converted and non-CT-converted human reference genome (GRCh38) k-mers.

The effect of seed length on sensitivity and speed can be inferred from the distribution of hash bin sizes for different seed lengths (Figure 4). With CT-converted seeds, we observe a flatter and broader distribution of bin sizes in comparison with the corresponding distribution for non-CT-converted seeds of equal length. This is reasonable, since CT-converted seeds ("3-letter alphabet") contain less information than non-CT-converted seeds ("4-letter alphabet"). In this way, one can see that the distribution of hash bin sizes for 25 nt CT-converted seeds approximates the distribution for 20 nt non-CT-converted seeds.

### Performance characteristics

In practice, the tradeoff between sensitivity and speed characterizes the performance of a read aligner. By adjusting runtime parameters (in particular, seed width, number of seeds examined per read, and maximum number of reference locations per seed), Arioc can be "tuned" to report additional mappings at the expense of speed. The shape of the speed-versus-sensitivity curves we observed illustrates that Arioc, like all read aligners, achieves increased sensitivity by spending additional computing time exploring a proportionally larger number of reference-sequence locations.

For read sequences that differ only minimally from the reference genome, Arioc uses search-space heuristics that cause it to find high-scoring mappings rapidly within a relatively small search space. For reads with multiple differences from the reference genome, Arioc examines more seed locations and computes more dynamic programming problems before it can report a satisfactory mapping. The consequent decrease in overall speed can be seen with "hard-to-align" sequencer reads in which a higher proportion of mappings have multiple differences from the reference (compare Supplementary Figure S9 with Figure 3).

Furthermore, Arioc (like any short-read aligner) explores a significantly larger search space when aligning BS-seq reads than it does when aligning non-bisulfite-treated reads. This has a tangible impact on throughput, which is typically about 75% of that observed for non-CT-converted reads.

### Implementation complexity and inaccuracies

The programming logic for handling bisulfite-treated reads is integrated into Arioc's software pipeline. This implementation avoids the software complexity and potential inaccuracies associated with BS-seq aligners that do not use this approach to the computation of BS-seq read alignments.

#### Inaccuracies due to wrapper implementation

The fundamental challenge in BS-seq alignment is to resolve the ambiguous representation of C_m_ (methylcytosine) in the original DNA as T in the sequencer reads. The straightforward approach used by a number of BS-seq aligners, including Bismark and GPU-BSM, is to compute alignments by comparing read sequences with a CT-converted reference sequence (genome).

This strategy is advantageous in that it can exploit an existing, well-optimized short-read aligner such as Bowtie 2 (Langmead and Salzberg, 2012) or SOAP3-DP (Luo et al., 2013) to carry out alignments. The BS-seq aligner implementation thus becomes a "wrapper" around an existing short-read aligner.

Although the benefit of such software reuse is immediately apparent, the design and implementation of such a wrapper implementation is complex. For example, to map a read sequence to the reverse complement of a reference genome, Bowtie 2 maps the reverse complement of the read sequence to the (forward) reference sequence. This technique obviously cannot work with a CT-converted reference sequence; to find reverse-complement mappings, the wrapper implementation must invoke Bowtie 2 a second time, using a GA-converted reverse-complement read sequence and a GA-converted index. In effect, the wrapper implementation aggregates mappings for each BS-seq read from two independent invocations of the short-read aligner. For reads in a non-directional library (in which each read sequence is represented by both its forward and reverse complement sequence), four such invocations are required (Krueger and Andrews, 2011).

Although the overhead of launching multiple read-aligner instances can be amortized to some extent by concurrent execution, the use of a general-purpose short-read aligner to compute alignments with CT-and GA-converted reads requires additional post-alignment processing before alignment results can be reported. The complex string-manipulation operations involved in post-alignment processing represent a performance bottleneck, especially for a wrapper implementation written in an interpreted language such as Python.

Post-alignment processing also introduces new sources of potential inaccuracy. In particular, the mappings produced by the wrapped short-read aligner cannot resolve the ambiguity inherent in CT conversion of the read sequences. Doing so requires the wrapper to identify occurrences of C_m_ through base-by-base comparison of each mapping with the reference sequence, re-scoring the alignment, and recomputing the CIGAR, MD, and MAPQ output for each reported read.

The wrapper implementation must also incorporate a method for choosing the "best" mapping(s) among those reported by different read-aligner instances. The usual technique is to report a mapping only when it is the unique, highest-scoring mapping for a read. This heuristic seemingly ensures that the implementation reports the "best" mapping (or no mapping at all) for each read, but it introduces reporting inaccuracies: the heuristic reports "best" mappings that have alignment scores that are only incrementally higher than other mapping(s) with the second-best alignment score, and filters out reads with equally high-scoring mappings in multiple regions of interest in the reference genome. The result is to blunt the overall sensitivity and specificity of the implementation. The magnitude of the problem is difficult to quantify but may be inferred from the shape and position of the ROC-like plots in Figure 2 and Supplementary Figures S1-S8.

#### Inaccuracies due to CT ambiguity

A further inaccuracy stems from the asymmetry of the CT ambiguity. When the read aligner identifies mappings using CT-converted sequences, it maps and scores C in the reference against T in the read sequence (indicating a bisulfite-converted, unmethylated cytosine) in the same way as T in the reference against C in the read sequence (indicating C_m_ in the read and a true mismatch at that base position). This situation is unusual, but when it occurs, the mappings for a read may be incorrectly scored and reported (Xi et al., 2012). It is possible, of course, to re-examine every mapping to detect this inaccuracy, but such additional post-processing would be costly in terms of overall throughput.

In any event, experience with BS-seq alignment has shown that these inaccuracies do not represent insurmountable obstacles to the use of a wrapper implementation. Nevertheless, these problems can be avoided within the read aligner by computing alignments directly between the original bisulfite-treated read sequences and the original reference sequence. Arioc encapsulates this logic within the software modules that compute and score read alignments, so the additional code required to align BS-seq reads executes on GPU threads and concurrent CPU threads.

### How fast is Arioc?

Our results imply that Arioc's BS-seq implementation is an order of magnitude faster than the most widely used non-GPU aligners. Of course, this estimate depends on GPU and CPU clock speeds, the number of available GPU and CPU threads, whether the aligner is parameterized to favor speed or sensitivity, and (for large data sets) disk I/O bandwidth.

Significantly, the speed results we report here are conservative because we want them to be directly comparable with results we previously reported for general short-read alignment. We therefore used the same Nvidia K20c devices (Nvidia, 2012) for both evaluations; with newer GPUs with higher internal clock speeds and more on-device memory, throughput is increased by an additional 40% for equivalent software parameterization settings.

Overall, Arioc provides a tangible increase in throughput in comparison with CPU-based BS-seq aligner implementations while maintaining high sensitivity and avoiding the most common potential inaccuracies associated with BS-seq read-alignment software. Arioc's speed also increases appropriately when additional CPU threads and multiple GPU devices are available.

These characteristics make Arioc a reasonable choice for aligning large datasets of bisulfite-treated short reads.

## ACKNOWLEDGEMENTS

We are grateful to Andrea Manconi and to Felix Krueger for their help with software configuration and for their insights into the technical challenges of BS-seq alignment.

## Supplementary Results

### Tables

Table T1 Candidates for performance comparisons.

Table T2 Software versions.

Table T3 Software configuration parameters.

Table T4 Distance between simulated and reported mapping positions.

### Results for simulated unpaired reads

Figure S1 Correctly mapped versus incorrectly mapped reads, for 100,000 simulated 100nt unpaired Illumina reads. Empirical error rate 0%, directional.

Figure S2 Correctly mapped versus incorrectly mapped reads, for 100,000 simulated 100nt unpaired Illumina reads. Empirical error rate 0%, non-directional.

Figure S3 Correctly mapped versus incorrectly mapped reads, for 100,000 simulated 100nt unpaired Illumina reads. Empirical error rate 5%, directional.

Figure S4 Correctly mapped versus incorrectly mapped reads, for 100,000 simulated 100nt unpaired Illumina reads. Empirical error rate 5%, non-directional.

### Results for simulated paired-end reads

Figure S5 Correctly mapped versus incorrectly mapped reads, for 100,000 simulated 100nt paired-end Illumina reads (200,000 mates). Empirical error rate 0%, directional.

Figure S6 Correctly mapped versus incorrectly mapped reads, for 100,000 simulated 100nt paired-end Illumina reads (200,000 mates). Empirical error rate 0%, non-directional.

Figure S7 Correctly mapped versus incorrectly mapped reads, for 100,000 simulated 100nt paired-end Illumina reads (200,000 mates). Empirical error rate 5%, directional.

Figure S8 Correctly mapped versus incorrectly mapped reads, for 100,000 simulated 100nt paired-end Illumina reads (200,000 mates). Empirical error rate 5%, non-directional.

### Results for BS-seq reads

Figure S9 Speed (reads aligned per second) versus sensitivity (percentage of paired-end reads mapped) for three BS-seq aligners, for 4 million 100 nt paired-end reads with a high proportion of low-scoring alignments.

Figure S10 Throughput (reads aligned per second) using one, two, and three GPUs (NVidia K20c) in a single computer for the data shown in Figure 3.

**Table T1.**
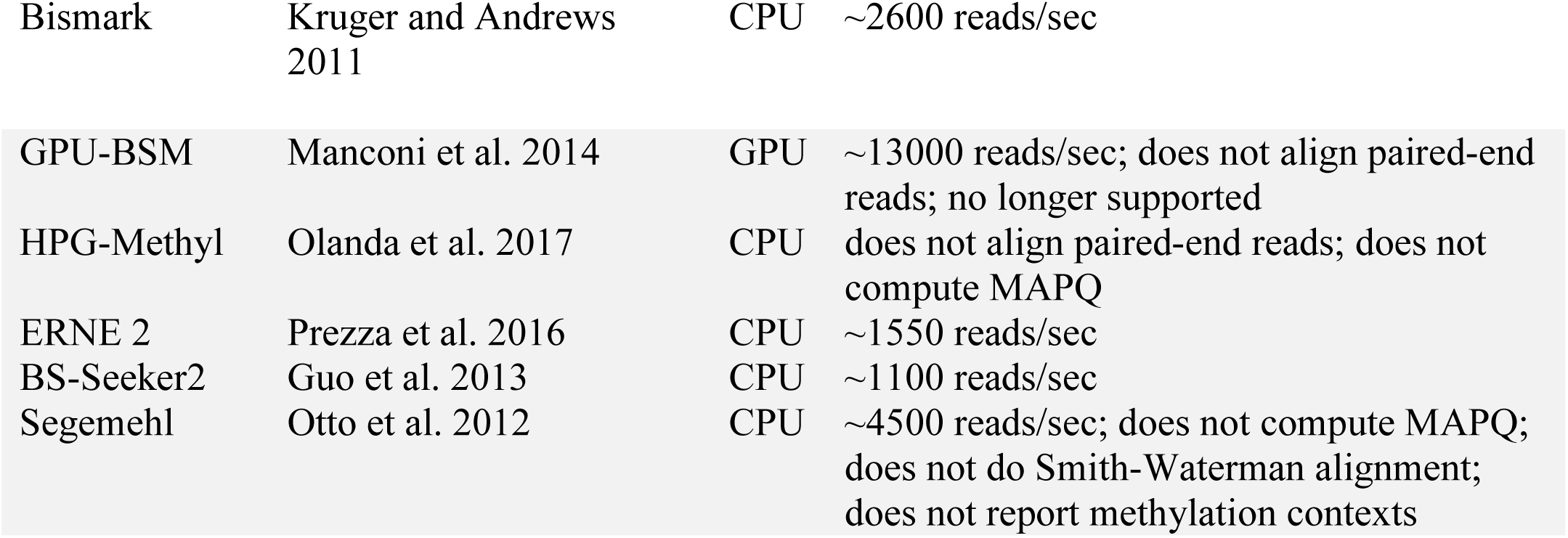
Candidates for performance comparisons. We considered the above CPU-based and GPU-based read aligners for detailed speed and sensitivity comparisons. Speed estimates are derived from published data. We carried out complete performance comparisons only with aligners that can handle both unpaired and paired-end mapping and that are capable of computing alignments on a large number (hundreds of millions) of short (100nt-250nt) reads on a single computer. We excluded aligners whose claimed or reported speed was not at least that of Bismark, or for which practical considerations (lack of support for all SAM/BAM fields, Smith-Waterman scoring, and methylation context) precluded a direct comparison using both simulated and sequencer-generated datasets.

|  |  |  |  |
| --- | --- | --- | --- |
| Bismark | Kruger and Andrews 2011 | CPU | ~2600 reads/sec |
| GPU-BSM | Manconi et al. 2014 | GPU | ~13000 reads/sec; does not align paired-end reads; no longer supported |
| HPG-Methyl | Olanda et al. 2017 | CPU | does not align paired-end reads; does not compute MAPQ |
| ERNE 2 | Prezza et al. 2016 | CPU | ~1550 reads/sec |
| BS-Seeker2 | Guo et al. 2013 | CPU | ~1100 reads/sec |
| Segemehl | Otto et al. 2012 | CPU | ~4500 reads/sec; does not compute MAPQ; does not do Smith-Waterman alignment; does not report methylation contexts |

---

**Table T2.** Software versions. All binaries executed using Red Hat Scientific Linux release 7.3 and NVidia CUDA v7.5.

|  |  |
| --- | --- |
| Arioc | 1.21 |
| Bismark | 0.18.1 |
| GPU-BSM | 2.7.3 |
| Segemehl | 0.2.0 (revision 418) |

**Table T3.** Software configuration parameters. Non-default parameters for the two extreme data points in Figure 3 (speed versus sensitivity). Arioc and Bismark were configured to perform local alignment using 25nt seeds. For Arioc, maxJ specifies the maximum size of a "bucket" in the seed-and-extend hash table; seedDepth limits the number of seed iterations.

|  | High throughput | High sensitivity |
| --- | --- | --- |
| Arioc | `<gapped maxJ="3" seedDepth="3" />` | `<gapped maxJ="200" seedDepth="3" />` |
| Bismark | `--p 6` | `--p 6` |
| Segemehl | `-D 0 -t 16` | `-D 1 -t 16` |

**Table T4.** Number of reads with distance D between simulated and mapped positions reported by Arioc for 100,000 simulated 100nt unpaired and 100,000 paired-end Illumina reads, at 0% and 5% error rate. For each mapping reported by each aligner, we used the range of reference-sequence positions reported by Sherman (in the FASTQ defline associated with each read) to compute a distance metric D that represented the smallest distance from each of the endpoints of the mapping to each of the endpoints of the range. Based on the above data, we chose a distance of 40 as a threshold for determining whether a simulated read was correctly mapped.

| **D** | **unpaired** |  | **paired-end** |  |
| --- | --- | --- | --- | --- |
|  | **0%** | **5%** | **0%** | **5%** |
| 0 | 96789 | 48701 | 98643 | 50516 |
| 1 | 1 | 1012 | 98940 | 52546 |
| 2 | 0 | 1554 | 87 | 2572 |
| 3 | 0 | 2828 | 82 | 4430 |
| 4 | 0 | 5717 | 47 | 8716 |
| 5 | 5 | 7063 | 250 | 13232 |
| 6 | 1 | 7871 | 397 | 15636 |
| 7 | 2 | 5889 | 340 | 14560 |
| 8 | 0 | 4953 | 234 | 11165 |
| 9 | 0 | 3712 | 224 | 9231 |
| 10 to 19 | 3 | 6669 | 344 | 18020 |
| 20 to 29 | 4 | 26 | 44 | 69 |
| 30 to 39 | 12 | 25 | 43 | 39 |
| 40 to 49 | 2 | 7 | 44 | 34 |
| 50 to 59 | 3 | 15 | 39 | 21 |
| 60 to 69 | 2 | 4 | 26 | 24 |
| 70 to 79 | 3 | 6 | 26 | 22 |
| 80 to 89 | 1 | 5 | 20 | 15 |
| 90 to 99 | 3 | 8 | 24 | 13 |
| 100+ | 11487 | 20321 | 31767 | 26733 |

**Figure S1.**
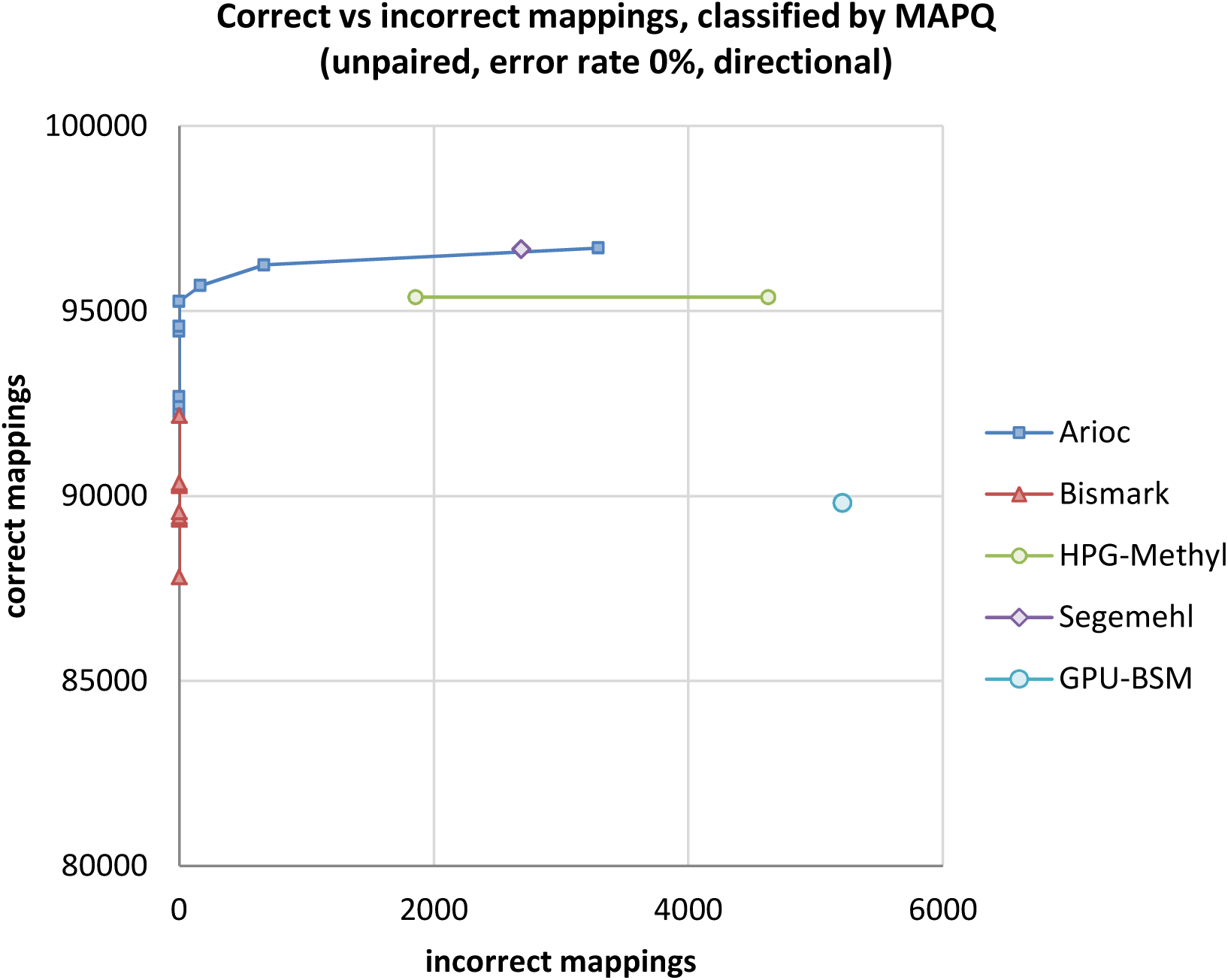
Total correctly mapped versus incorrectly mapped reads, plotted for decreasing MAPQ, for 100,000 simulated 100nt unpaired Illumina reads. Empirical error rate: 0% (Sherman parameters: -cr 50.0)

**Figure S2.**
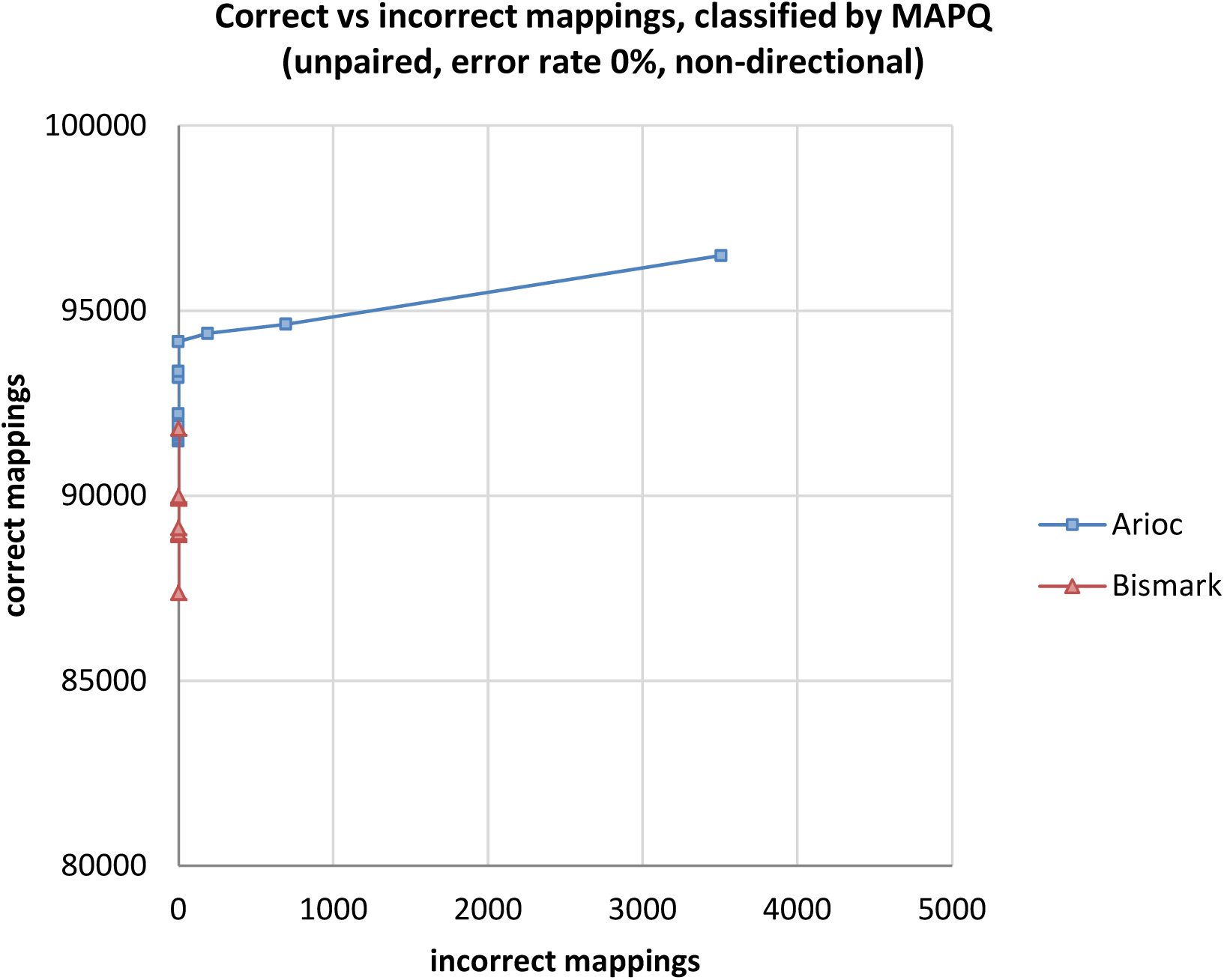
Total correctly mapped versus incorrectly mapped reads, plotted for decreasing MAPQ, for 100,000 simulated 100nt unpaired Illumina reads. Empirical error rate 0% (Sherman parameters: -cr 50.0 --non_directional).

**Figure S3.**
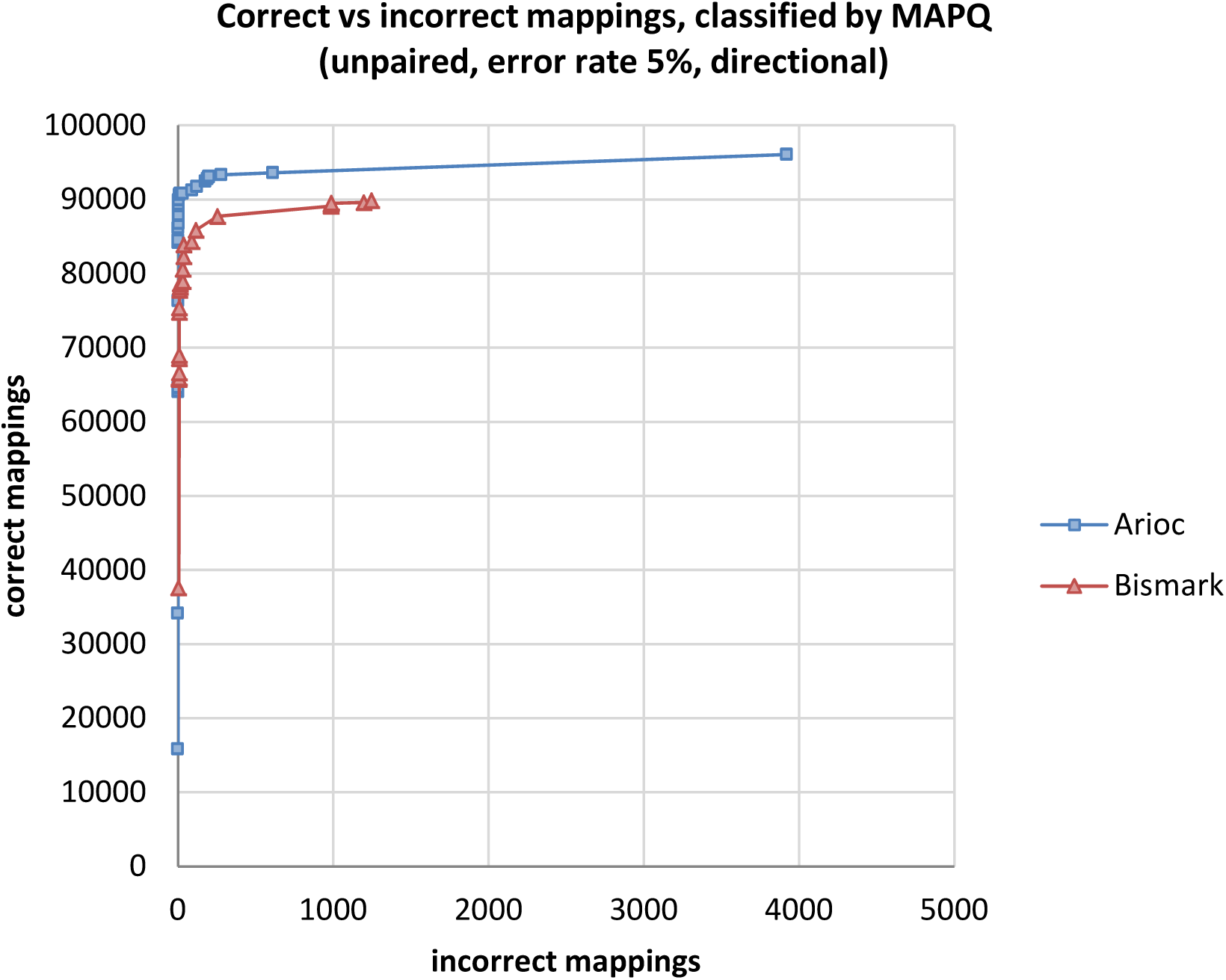
Total correctly mapped versus incorrectly mapped reads, plotted for decreasing MAPQ, for 100,000 simulated 100nt unpaired Illumina reads. Empirical error rate 5% (Sherman parameters: -cr 50.0 -e 5.0).

**Figure S4.**
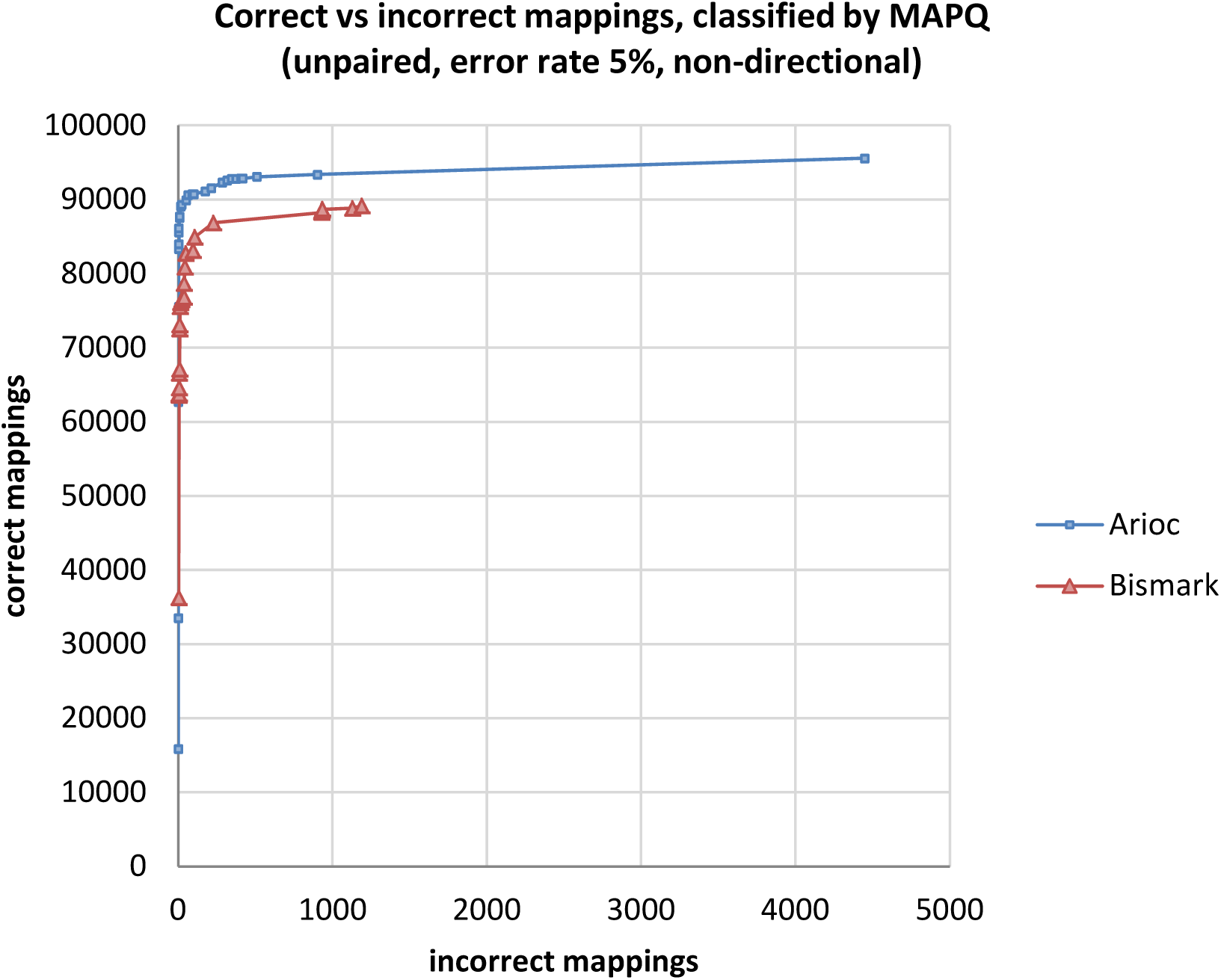
Total correctly mapped versus incorrectly mapped reads, plotted for decreasing MAPQ, for 100,000 simulated 100nt unpaired Illumina reads. Empirical error rate 5% (Sherman parameters: -cr 50.0 -e 5.0 --non_directional).

**Figure S5.**
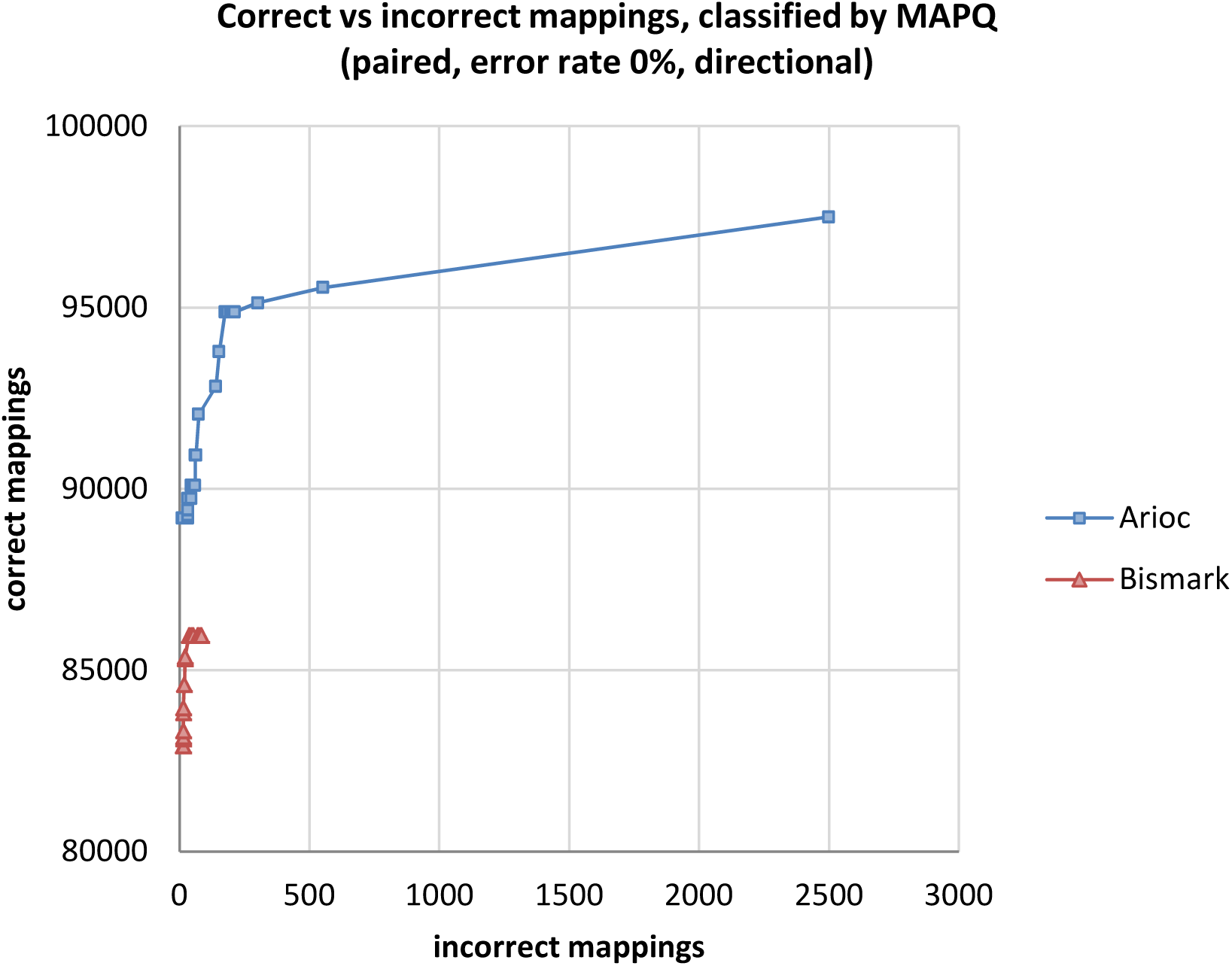
Total correctly mapped versus incorrectly mapped reads, plotted for decreasing MAPQ, for 100,000 simulated 100nt paired-end Illumina reads (200,000 mates). Empirical error rate: 0% (Sherman parameters: -pe -cr 50.0) (Same as Figure ???.)

**Figure S6.**
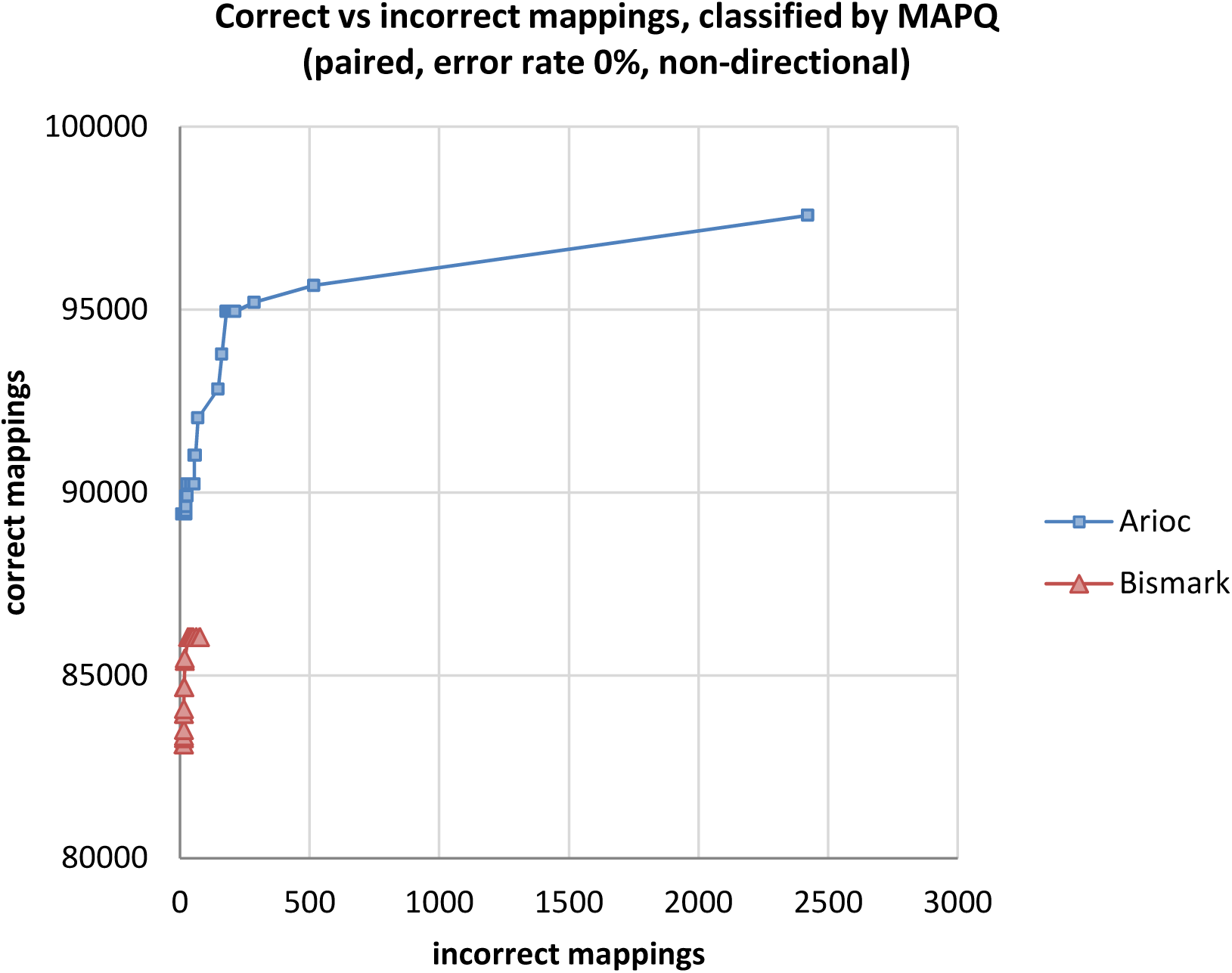
Total correctly mapped versus incorrectly mapped reads, plotted for decreasing MAPQ, for 100,000 simulated 100nt paired-end Illumina reads (200,000 mates). Empirical error rate 0% (Sherman parameters: -pe -cr 50.0 --non_directional).

**Figure S7.**
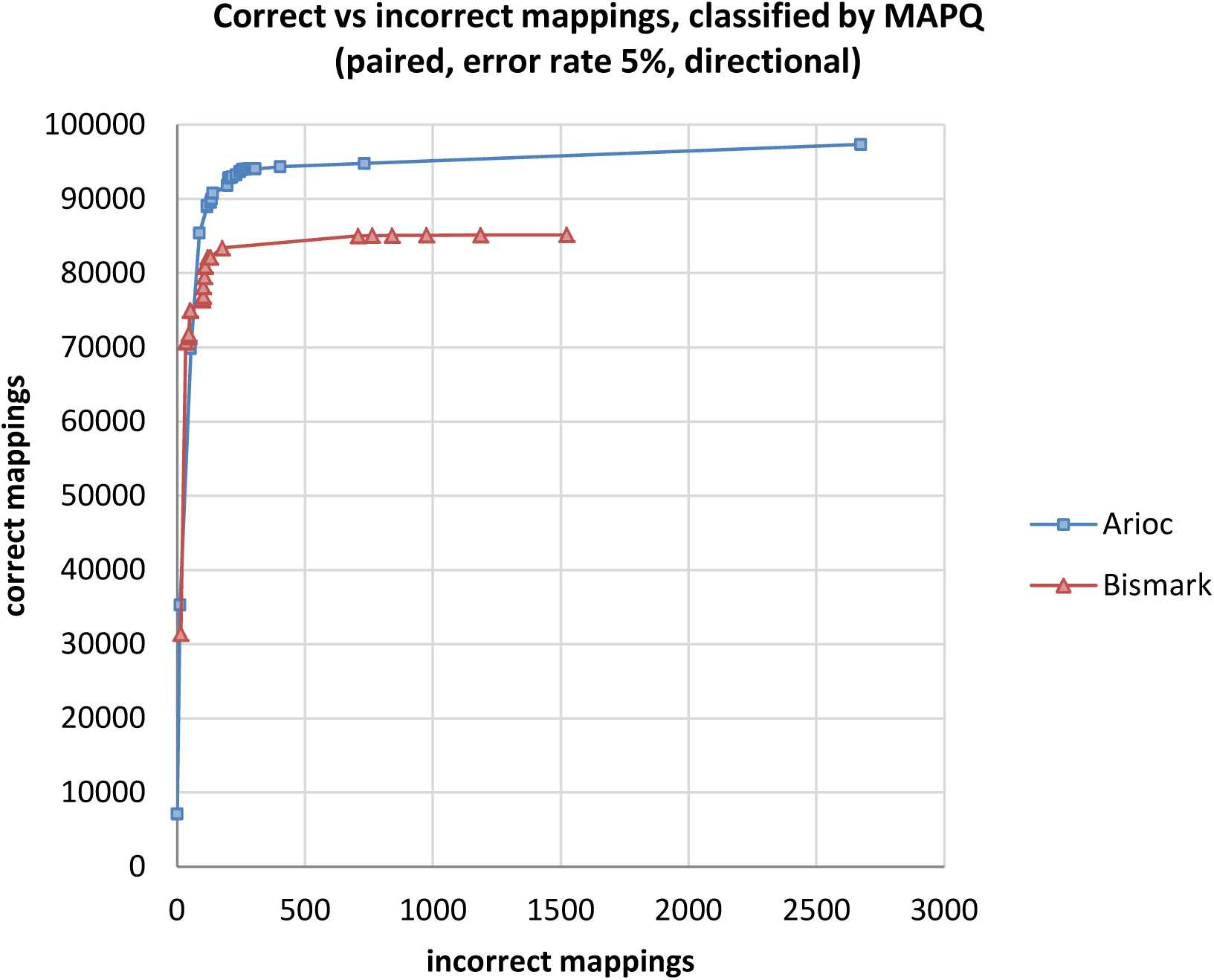
Total correctly mapped versus incorrectly mapped reads, plotted for decreasing MAPQ, for 100,000 simulated 100nt paired-end Illumina reads (200,000 mates). Empirical error rate 5% (Sherman parameters: -pe -cr 50.0 -e 5.0).

**Figure S8.**
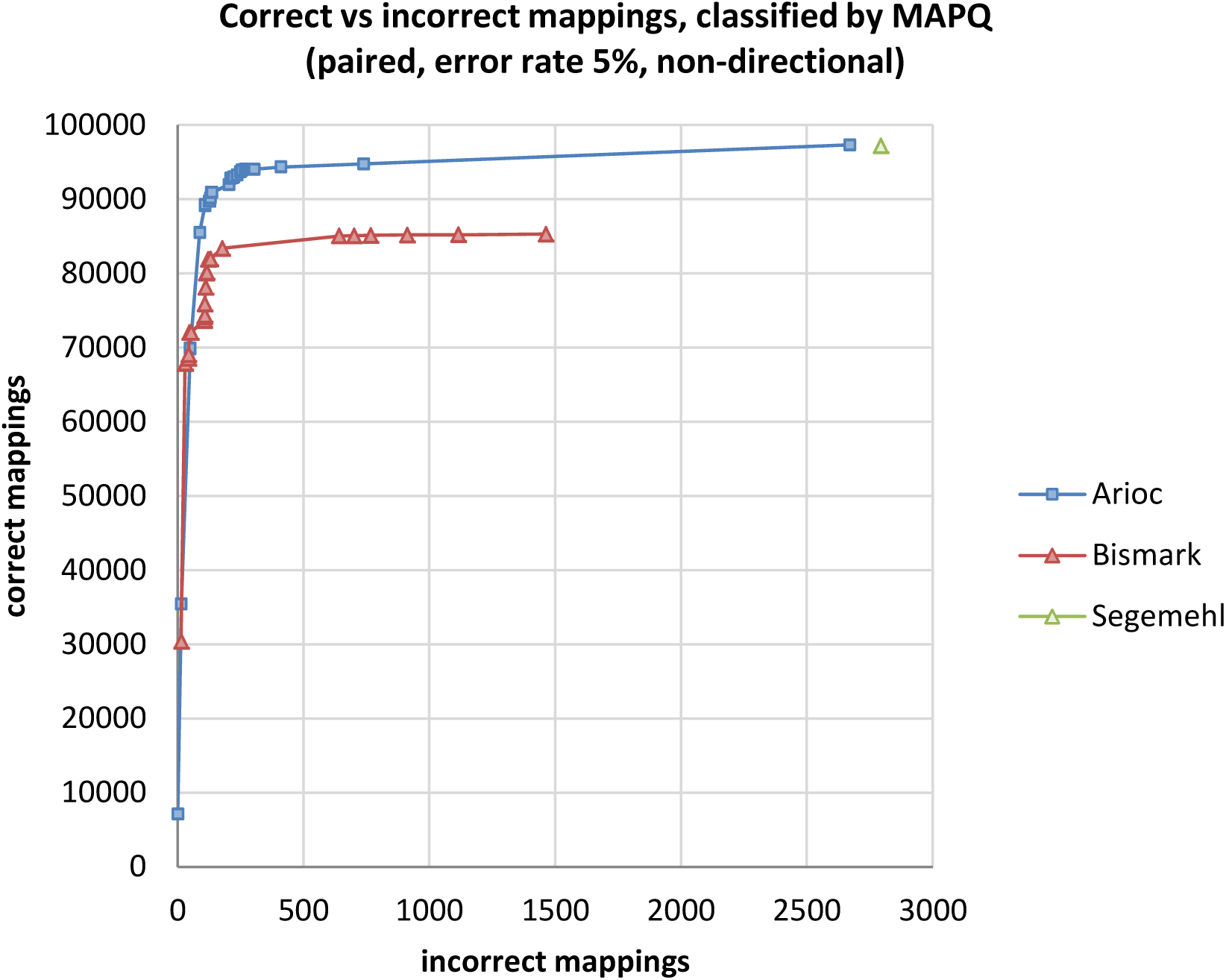
Total correctly mapped versus incorrectly mapped reads, plotted for decreasing MAPQ, for 100,000 simulated 100nt paired-end Illumina reads (200,000 mates). Empirical error rate 5% (Sherman parameters: -pe -cr 50.0 -e 5.0 --non_directional).

**Figure S9.**
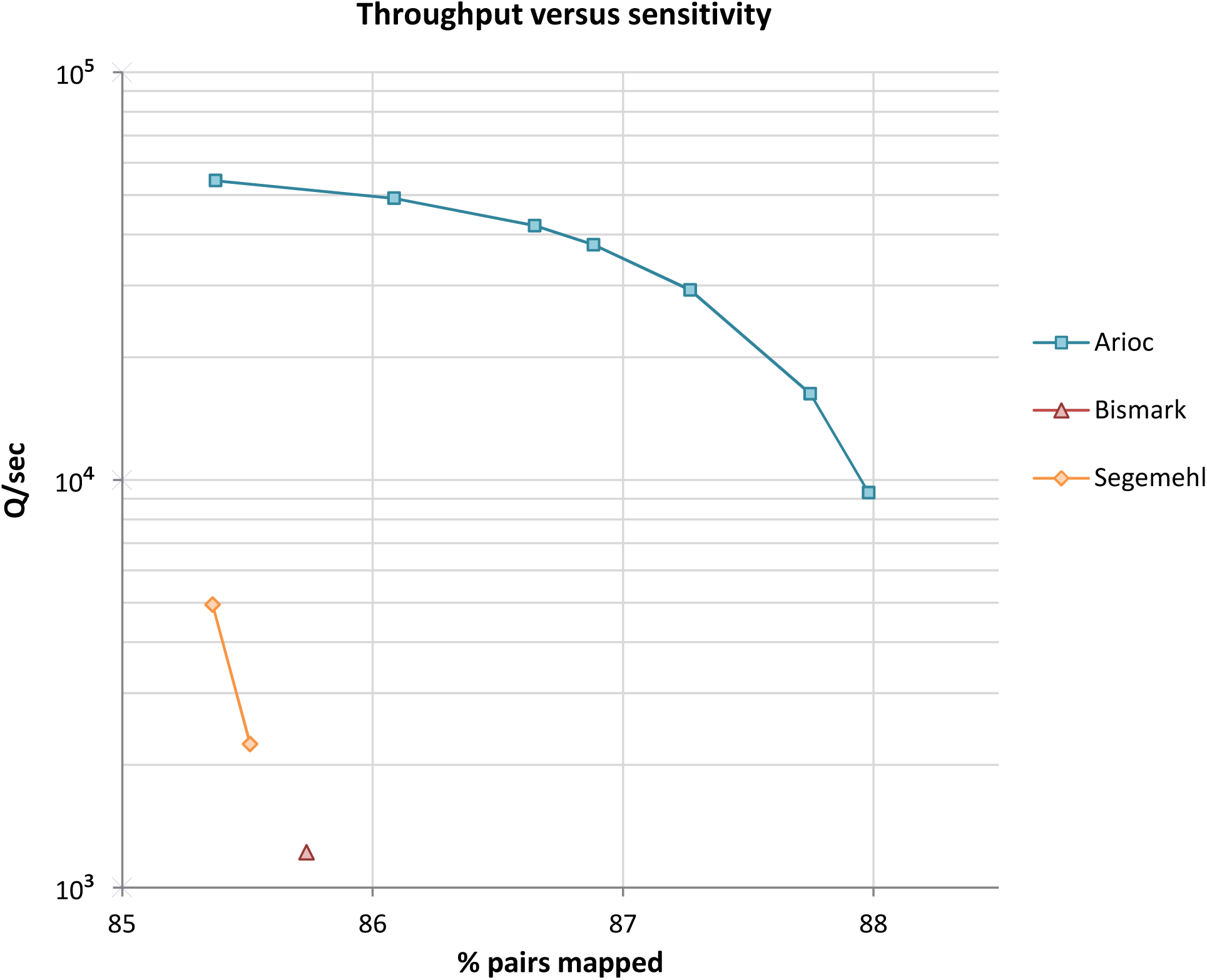
Speed (reads aligned per second) versus sensitivity (percentage of paired-end reads mapped) for three BS-seq aligners. Data for 4 million 100 nt paired-end reads (unpublished data) with a high proportion of low-scoring alignments. Workstation hardware: same as Figure 3.

**Figure S10.**
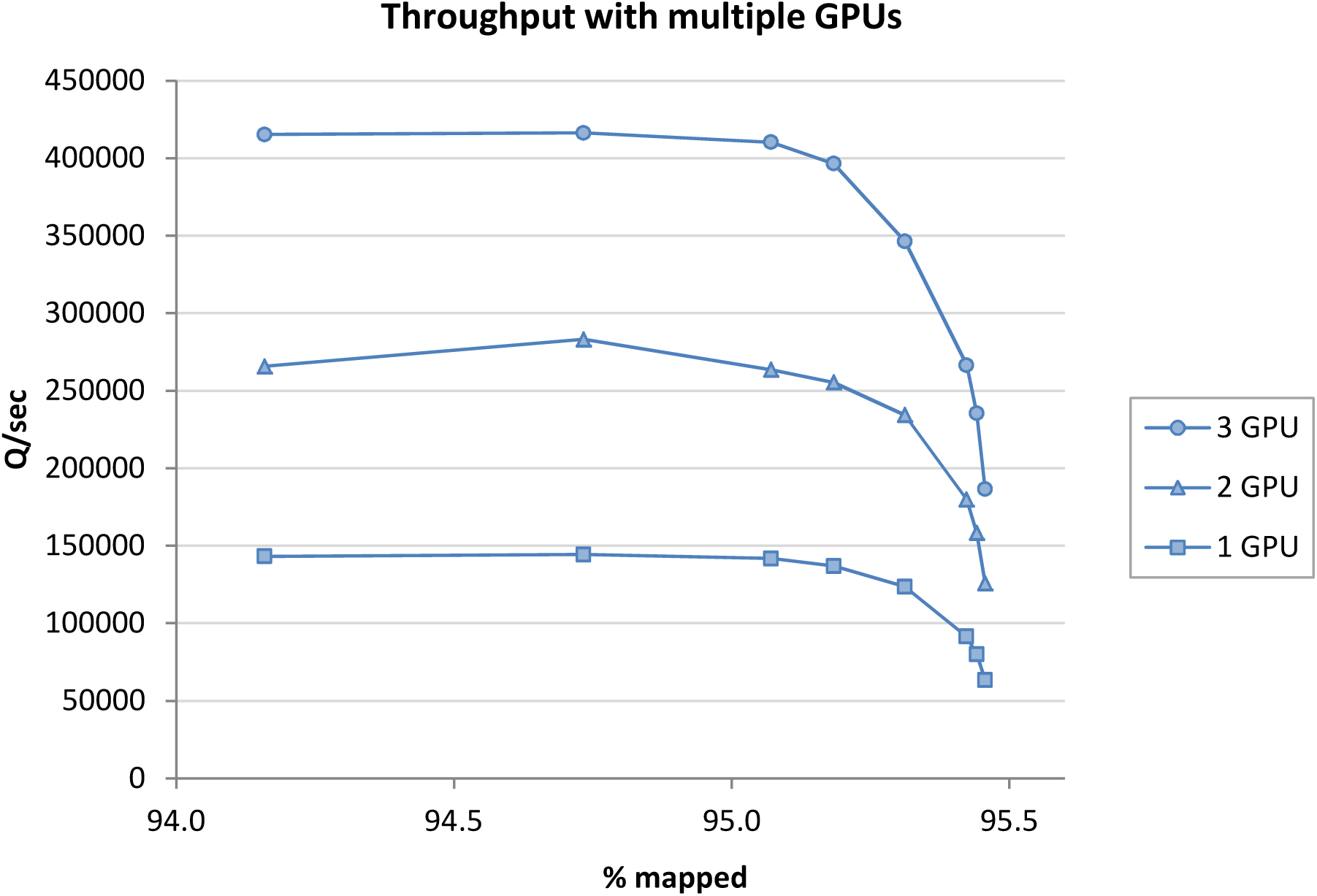
Throughput (reads aligned per second) using one, two, and three GPUs (NVidia K20c) in a single computer for the data shown in Figure 3.

